# The *Phtheirospermum japonicum* isopentenyltransferase PjIPT1a regulates host cytokinin responses in *Arabidopsis*

**DOI:** 10.1101/2021.05.24.445531

**Authors:** Anne Greifenhagen, Isabell Braunstein, Jens Pfannstiel, Satoko Yoshida, Ken Shirasu, Andreas Schaller, Thomas Spallek

## Abstract

The hemiparasitic plant *Phtheirospermum japonicum* is a nutritional specialist that supplements its nutrient requirements by parasitizing other plants through haustoria. During parasitism, the *Phtheirospermum* haustorium transfers hypertrophy-inducing cytokinins (CKs) to the infected host root. The CK biosynthesis genes required for haustorium-derived CKs and the induction of hypertrophy are still unknown. We searched for haustorium-expressed isopentenyltransferases (IPTs) that catalyse the first step of CK biosynthesis, confirmed the specific expression by *in vivo* imaging of a promoter-reporter, and further analysed the subcellular localization, the enzymatic function, and contribution to inducing hypertrophy by studying CRISPR-Cas9 induced *Phtheirospermum* mutants. *PjIPT1a* was expressed in intrusive cells of the haustorium close to the host vasculature. PjIPT1a and its closest homolog PjIPT1b located to the cytosol and showed isopentenyltransferases activity *in vitro* with differences in substrate specificity. Mutating *PjIPT1a* abolished parasite-induced CK responses in the host. A homolog of *PjIPT1a* with shared characteristics was also identified in the related weed *Striga hermonthica*. We propose that *PjIPT1a* exemplifies how parasitism-related functions evolve through gene duplications and neofunctionalization.

## Introduction

Most parasitic species maintain some level of photosynthesis (hemiparasites), but access soil nutrients mainly by parasitizing other plants (Heide-Jorgensen, 2008; Brundrett & Tedersoo, 2018). The two main lineages of hemiparasitic plants, Santalales and the family of Orobanchaceae in the Lamiales account for more than 90% of all parasitic species (Nickrent, 2020). The greatest in-family diversity of hemiparasitic plants is found in the Orobanchaceae. In contrast to the agriculturally important weed *Striga hermonthica*, non-weedy species such as the model parasitic plant *Phtheirospermum japonicum* can propagate without being attached to a host (Clarke *et al.*, 2019).

Genetic adaptations to the parasitic lifestyle start to be revealed from the sequencing of transcriptomes and genomes of parasitic plants (Yang *et al.*, 2015; Vogel *et al.*, 2018; Sun *et al.*, 2018; Yoshida *et al.*, 2019). At the centre of these studies is the haustorium, a multi-cellular organ that facilitates infection and parasitism (Yoshida *et al.*, 2016; Kokla & Melnyk, 2018). Comparative transcriptomics in three Orobanchaceae species with varying degrees of host dependencies unearthed remarkable conservation of the transcriptional signatures during infection (Yang *et al.*, 2015). Further studies confirmed these tendencies also in other lineages (Vogel *et al.*, 2018; Sun *et al.*, 2018; Yoshida *et al.*, 2019). The term “core parasitism genes” describes the shared transcriptional behaviour of a group of homologous genes within a parasitic plant lineage (Yang *et al.*, 2015).

The concept of core parasitism genes may be extended to “core parasitism strategies”. Strategies that are commonly observed during plant parasitism include the secretion of cell wall degrading enzymes and changes in hormone homeostasis (Brun *et al.*, 2020). For example, cytokinins (CKs) are produced in haustoria of *Cuscuta japonicum*, *Santalum album* and *Phtheirospermum japonicum* (Zhang *et al.*, 2012; Furuhashi *et al.*, 2013; Spallek *et al.*, 2017). Transfer of cytokinin from the parasite to the host was demonstrated using *Arabidopsis* cytokinin biosynthesis mutants and transgenic *Phtheirospermum* roots that express a cytokinin-degrading enzyme (Spallek *et al.*, 2017). With prolonged cytokinin transfer, infected host tissue becomes hypertrophic, a symptom that is also induced by mistletoes, *Viscum album* (Loranthaceae in the order Santalales), and *Alectra vogelii* (Orobanchaceae) (Heide-Jorgensen, 2008).

Cytokinins are *N^6^*-substituted adenine derivatives, of which isoprenoid cytokinins are most prevalent in plants (Sakakibara, 2006). *N^6^*-prenylation is catalysed by adenosine phosphate-isopentenyltransferases (IPTs, Fig. **1a**). Plant IPTs prefer dimethylallyl diphosphate (DMAPP) as isoprenoid donor, and ATP and ADP (ATP/ADP-type IPTs) or tRNA (tRNA-type IPTs) as isoprenoid acceptors (Kakimoto, 2001; Sakakibara, 2006). The agrobacterial IPTs, Tmr and Tzs, prefer hydroxy-methylbutenyl diphosphate and AMP as substrates. Tmr and Tzs are required for virulence and crown-gall formation (Akiyoshi *et al.*, 1984; Heinemeyer *et al.*, 1987). Overexpression of *IPT*s in *Arabidopsis* causes ectopic cytokinin responses (Kakimoto, 2001) indicating that IPTs catalyse the rate limiting step in cytokinin biosynthesis. In *Arabidopsis*, the major route of cytokinin biosynthesis relies on ATP/ADP-type IPTs targeted to plastids (AtIPT1, AtIPT3, AtIPT5, AtIPT8), to mitochondria (AtIPT7), or residing in the cytosol and nucleus (AtIPT3, AtIPT4). tRNA-type IPTs (AtIPT2, AtIPT9) also localize to the cytosol and nucleus (Kasahara *et al.*, 2004; Kieber & Schaller, 2014). The biological relevance of different subcellular IPT locations is not known.

**Figure 1:**
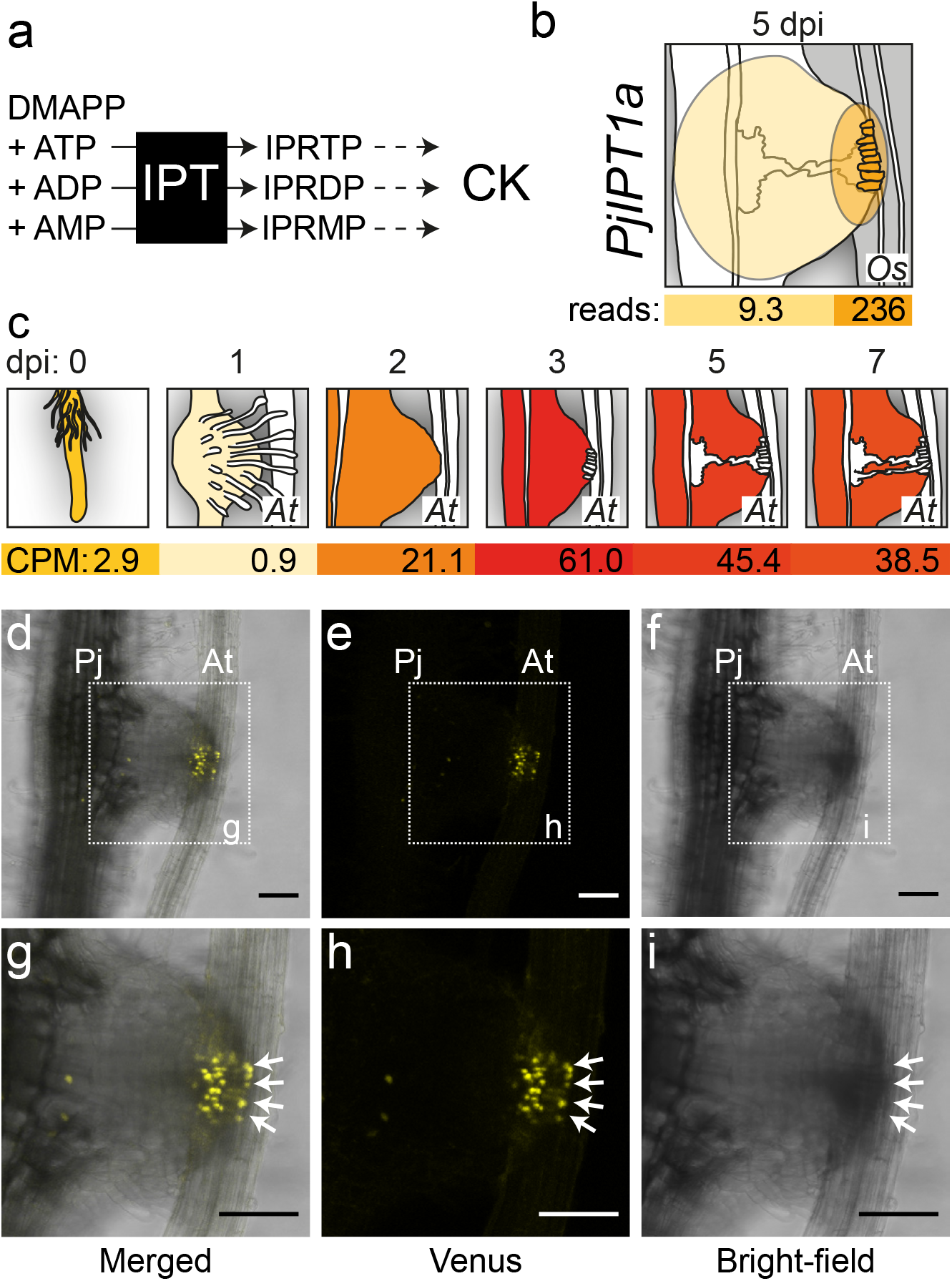
*PjIPT1a* is specifically expressed in intrusive cells. (a) Prenylation of ATP, ADP and AMP by isopentenyltransferases (IPT) with di-methylallyl pyrophosphate (DMAPP) to isopentenyladenosine-5’-triphosphate (IPRTP), isopentenyladenosine-5’-diphosphate (IPRDP) and isopentenyladenosine-5’-monophosphate (IPRMP) is the first step of cytokinin (CK) biosynthesis. (b) RNA-Sequencing data (originally published in Ogawa et al., 2020) shows increased expression of *PjIPT1a* in intrusive cells of *Phtheirospermum* haustoria 5 days post-infection (dpi) of rice roots (*Oryza sativa*, *Os*). (c) Upregulation of *PjIPT1a* expression during infection of *Arabidopsis* (*At*) roots peaks with 61 CPM (counts per million mapped reads) at 3 dpi (RNA-Sequencing data from Kurotani et al., 2020). (b, c) Colour gradients (white-yellow-red) relate to *PjIPT1a* expression in indicated sampling areas. Haustoria diagrams are based on haustoria outlines presented in Brun et al. 2020. (d - i) Confocal images show a *Phtheirospermum* (*Pj*) haustorium expressing *pIPT1:3xVenus-NLS* during infection of *Arabidopsis* (*At*) at 4 dpi. Merged yellow channel for Venus detection and bright-field is depicted in (d), individual Venus and bright-field channels in (e) and (f), respectively. Zoomed-in scans shown in g - i correspond to the region marked by dotted lines in (d – f). Intrusive cells are indicated with white arrows. Scale bars = 100 μm.

Here, we report the function of *Phtheirospermum* PjIPT1a as essential for triggering *Phtheirospermum*-induced cytokinin responses in *Arabidopsis.* PjIPT1a is an unconventional plant isopentenyltransferase with parasitism-related features. PjIPT1a uses AMP and ATP as prenyl acceptors and is located in the cytosol. It is expressed in intrusive cells of the *Phtheirospermum* haustorium. Targeted gene-knockouts of *PjIPT1a* by CRISPR/Cas9 suppressed parasite-induced cytokinin responses in the host. Similarities to isopentenyltransferases from *Striga* suggest that hemiparasitic Orobanchaceae use common strategies to manipulate host plants.

## Materials and Methods

### Plant growth conditions and transformation

*Phtheirospermum japonicum* (Okayama, wild-type) seeds were sterilized in 70% ethanol, 0.05% SDS for 10 minutes, and then washed with 98% ethanol. For transformation, plants were grown on Gamborg B5 medium (Duchefa) supplemented with 1% sucrose at 25°C under long-day conditions (16 h light). Seven-days-old *Phtheirospermum* seedlings were transformed with *Agrobacterium rhizogenes* AR1193 as previously described (Ishida *et al.*, 2011). Approximately four weeks after transformation, *Phtheirospermum* seedlings with hairy roots were transferred to 0.8% agar containing 300 μg/ml cefotaxime. Two days later, *Arabidopsis* Ws *pARR5:GFP* seedlings grown for one week on MS (Duchefa) plates were aligned next to *Phtheirospermum* hairy roots. One week later, infected plants were transferred to half-strength MS plates and grown for additional 21 days. *Nicotiana benthamiana* plants were grown in soil under long-day conditions at 25°C.

### Phylogenetic analysis

IPT amino acid sequences were retrieved from public databases (NCBI), and by blast search using IPT protein sequences as query against published genomes (Table **S1**). CLC Main Workbench Version 8 was used to align protein sequences and to construct a phylogenetic tree based on the UPGMA and Jake-Cantor algorithm with a bootstrap analysis of 1000 replicates.

### Cloning, plasmid design and genotyping

We cloned the 2.6 kb upstream sequence of the *PjIPT1a* gene while removing an internal Bpi1site into a Golden Gate vector that was then used to generate 3xVenus-Nuclear Localization Signal (NLS)-based reporter as previously described (Engler *et al.*, 2014; Cui *et al.*, 2016). Primer sequences are given in Table **S2**. Open reading frames (ORFs) of *PjIPT1a* and *PjIPT1b* were first cloned into pENTR/D-TOPO entry vector (Thermo Fisher Scientific) to generate GFP fusion constructs using the pGWB5 binary vector (Nakagawa *et al.*, 2007). The putative chloroplast transit peptide was cloned into pART7 using XhoI and ClaI (Thermo Fisher Scientific) and then into pART27 (Gleave, 1992). An empty pGREEN vector was used to express *p35S:GFP* (Hellens *et al.*, 2000). Golden Gate-compatible vectors carrying *PjIPT1a*-specific gRNAs, *Cas9*, and the transformation marker *p35S:DsRED* were assembled as previously described (Nekrasov *et al.*, 2013; Engler *et al.*, 2014). gRNAs were chosen based on their specificity to the intended target. Individual hairy roots were genotyped by amplifying a 1651 bp long amplicon that contained the 954 bp *PjIPT1a* gene. *PjIPT1a* alleles were cloned into pBluescript-II-SK(+) vectors for sequencing. For expression in *E. coli*, ORFs of *PjIPT1a*, *PjIPT1b*, and *GST* were fused with a C-terminal octa-His tag in the pET42 vector (Novagen).

### Expression of proteins in *N. benthamiana* and *E. coli*

*Agrobacterium tumefaciens* GV3101 were infiltrated in a 10 mM MES (pH 5.6) and 10 mM MgCl_2_ buffer into approximately five-week-old *N. benthamiana* leaves, which were imaged two days later. *PjIPT1a-His*, *PjIPT1b-His*, and *GST-His* were expressed in *E. coli* BL21 RIL cells and purified on NiNTA beads (Qiagen), then concentrated and desalted using Vivaspin 500 columns (Sartorius).

### Microscopy

Hairy roots were imaged in water using a Zeiss Axio Image Z.1 fluorescence microscope. *PjIPT1a* promoter activity was recorded with a Leica TCS SP5 II, the subcellular location of fluorescent proteins with a Zeiss LSM 700 confocal microscope.

### IPT-*in vitro* assays

IPT-*in vitro* assays were conducted as previously described (Brugière *et al.*, 2008). In brief, 8 μg of purified proteins were incubated with 1 mM DMAPP (Sigma-Aldrich) and 1 mM AMP or ATP for two hours at 30°C in 12.5 mM Tris–HCl (pH 7.5), 37.5 mM KCl, 5 mM MgCl_2_. The reaction was stopped at 95°C and methanol precipitated. The supernatant was analysed by UHPLC-MS/MS on a 1290 UHPLC system (Agilent) coupled to a Q-Exactive Plus Orbitrap mass spectrometer equipped with a heated electrospray ionization (HESI) source (Thermo Fisher Scientific). Analyte separation was achieved by a Phenomenex Luna Omega Polar C18 column. Mobile phase A was 0.2% formic acid in water, and mobile phase B 0.2% formic acid in acetonitrile. The HESI source was operated in positive ion mode, with a capillary voltage of 4.2 kV and an ion transfer capillary temperature of 360°C. Mass spectra were acquired in MS mode within the mass range of 150 to 800 m/z at a resolution of 70,000 FWHM using an Automatic Gain Control (AGC) target of 3 × 106 of and 100 ms maximum ion injection time (MIT). Data dependent MS/MS spectra in a mass range of 200 to 2000 m/z were generated for the five most abundant precursor ions with a resolution of 17,500 FWHM using an AGC target of 5 × 104, 64 ms MIT and a stepped collision energy of 15, 30 and 55. Xcalibur software version 4.4.16.14 (Thermo Fisher Scientific) was used for instrument control and data acquisition. Compound Discoverer software 3.2 (Thermo Fischer Scientific) was used for analysing raw instrument data (.raw files) and peak area calculation.

### Statistical analysis

The R packages Rmisc, readr, multcompView, ggplot2 and ggpubr were used for statistics (R Core Team, 2020).

## Results

### *PjIPT1a* is expressed in intrusive cells of *Phtheirospermum* haustoria

CK biosynthesis is tightly regulated by environmental and developmental cues and restricted to distinct tissues and cells. We thus reasoned that CK production in *Phtheirospermum* haustoria requires the haustorium-specific expression of at least one *IPT* gene. We surveyed published *Phtheirospermum* transcriptomes for infection-specific expression of *IPTs* and identified one, hereinafter referred to as *PjIPT1a*, that is upregulated in the course of infection and shows 25-fold higher expression in intrusive cells relative to the rest of the haustorium (Fig. **1b**) (Ogawa *et al.*, 2020). *PjIPT1a* expression increases until day three of the infection and continues at a relatively high level for the later time points (Fig. **1c**) (Kurotani *et al.*, 2020). This expression profile coincides with the induction of CK response markers in parasite and host, which become detectable from 3 - 4 dpi (days post-infection) and remain visible until at least 30 dpi (Spallek *et al.*, 2017). Using a 2.6 kb *PjIPT1a* promoter fragment to drive the expression of a *3xVenus-NLS* reporter, we confirmed the specific expression of the *PjIPT1a* gene in intrusive cells of *Phtheirospermum* haustoria at 4 dpi of *Arabidopsis* roots (Fig. **1d-f**). *PjIPT1a* is thus expressed in cells in close, likely direct, contact to the host vasculature (Fig. **1g-i**).

### *PjIPT1a* encodes a cytosolic IPT with homologs in *Striga hermonthica*

To validate the phylogenetic relationship of PjIPT1a to other *Phtheirospermum* IPTs, we compared the amino acid sequences of PjIPTs to known plant IPTs and two agrobacterial IPTs, Tzs and Tmr (Fig. **2a**). PjIPT1a clustered with other plant IPT1s, and was more similar to proteins from the related parasitic species *Striga hermonthica* (Orobanchaceae, Lamiales) and *Streptocarpus rexii* (Gesneriaceae, Lamiales) than to IPTs from more distantly related non-parasitic species such as common hop (*Humulus lupulus*, Cannabaceae, Rosales) and *Arabidopsis thaliana* (Brassicaceae, Brassicales). Two close homologs were detected for PjIPT1a and its *Striga* ortholog ShIPT1a. Like *PjIPT1a*, only *ShIPT1a* is induced upon infections but not *ShIPT1b* (Fig. **S1**).

**Figure 2:**
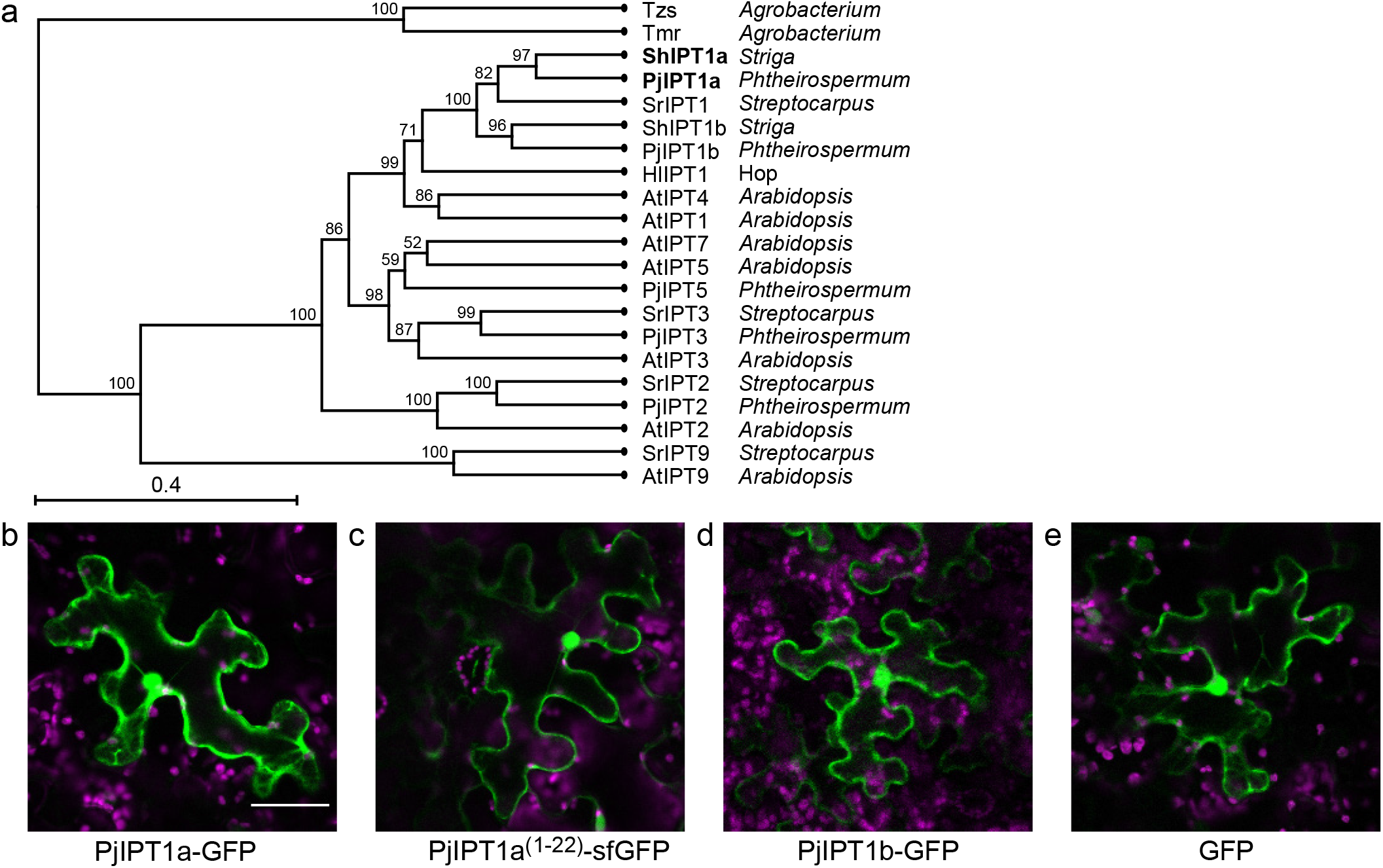
*PjIPT1a* and its closest homolog are nuclear-cytoplasmic IPTs. Phylogenetic analysis (UPGMA) based on the amino acid sequences of prokaryotic (*Agrobacterium*) and plant IPTs positions IPT1a within a cluster of plant IPT1s and closest to the *Striga* ShIPT1a protein. Bootstrap values are shown as percentage of 1000 repetitions. (b-e) PjIPT1a and PjIPT1b located in the cytoplasm and nucleus when expressed in *N. benthamiana*. The subcellular localization pattern of the full-length PjIPT1a-GFP (b) and PjIPT1b-GFP (d) fusions matched the pattern observed for PjIPT1a^(1-22)^-sfGFP (c) and free GFP (e). Scale bar = 50 μm.

Bioinformatics tools predict a chloroplast transit peptide (CTP) in the first 22 amino acids of PjIPT1a; however, at a lower score compared to AtIPT1 and SrIPT1 (Table **S3**). Putative CTPs of PjIPT1a and ShIPT1a are less than half as long as the CTPs predicted for AtIPT1 (71 amino acids) and SrIPT1 (46 amino acids) (Table **S3**; Fig. **S2**). Therefore, we experimentally tested subcellular localization of PjIPT1a and functionality of its CTP. A PjIPT1a-GFP fusion protein transiently expressed in *N. benthamiana* located to cytosol and nucleus (Fig. **2b**), with no apparent difference to the localization of free GFP (Fig. **2e**). Western-blots using anti-GFP antibodies confirmed the integrity of the PjIPT1a-GFP full-length protein (Fig. **S3**). The data thus suggest nuclear-cytoplasmic localization of PjIPT1a.

Nuclear-cytoplasmic localization is also observed for chloroplast-targeted AtIPT3, when it is post-translationally modified by C-terminal farnesylation (Galichet *et al.*, 2008). To exclude the possibility that post-translational modifications of PjIPT1a outside the putative CTP would affect its localization, we analysed the subcellular targeting of super-folder (sf) GFP fused to the predicted transit peptide (the first 22 amino acids). The localization of PjIPT1a^(1–22)^-sfGFP (Fig. **2c**) was indistinguishable from PjIPT1a-GFP and free GFP (Fig. **2b,e**). The same nuclear-cytoplasmic localization was also observed for PjIPT1b-GFP (Fig. **2d**). The putative CTP signal in *Striga* ShIPT1a is even shorter (11 amino acids) than the apparently non-functional CTP of PjIPT1a (Fig. **S2**; Table **S3**), while no CTP was predicted for ShIPT1b. Therefore, we assume a similar subcellular localization pattern of the homologous ShIPT1a and ShIPT1b proteins.

### PjIPT1a drives CKs responses in host plants

Haustorium-derived CKs induce hypertrophy of the host root and can be visualized by a host-expressed cytokinin reporter (*pARR5:GFP*) (Spallek *et al.*, 2017). To address a role for *PjIPT1a* in the production of haustorium-derived CKs, we generated transgenic *Phtheirospermum* roots (hairy roots, HR) that expressed the *Cas9* gene together with two *PjIPT1a*-specific gRNAs (Fig. **3b**). The construct also contained *p35S:dsRED* as a fluorescent marker for identification of transformed roots. HR transformed with *p35S:dsRED* alone (HR1) and hairy roots without the *Cas9/PjIPT1a-gRNA*s + *dsRED* cassette (HR2) served as controls. In contrast to control HRs (HR1-2) that triggered *pARR5:GFP* expression and hypertrophic growth in infected *Arabidopsis* roots, neither *pARR5:GFP* expression nor hypertrophic growth were observed after infection with HRs containing the *Cas9/PjIPT1a-gRNA*s + *dsRED* cassette (HR3-5) (Fig. **3a**). While the *PjIPT1a* locus was successfully PCR-amplified in full length from control HR1-2 (Fig. **3c**), additional smaller bands were observed for *Cas9*-containing HR3-5 (Fig. **3c**). Sequencing *PjIPT1a* amplicons revealed Cas9-induced deletions of the *PjIPT1a* locus ranging from single nucleotides to 1100 bp for HR3-5 (Fig. **3d**). For HR4, we identified three alleles that carried deletions of 40 bp, 893 bp, and 1110 bp in the *PjIPT1a* gene, indicating a chimeric pattern of Cas9-induced mutations in *Phtheirospermum* hairy roots (Fig. **3d**). The presence of Cas9-induced deletions in *PjIPT1a* correlated perfectly with the absence of a *Phtheirospermum*-induced cytokinin response in *Arabidopsis*. We therefore conclude that *PjIPT1a* is an essential component in the CK transfer from parasite to host.

**Figure 3:**
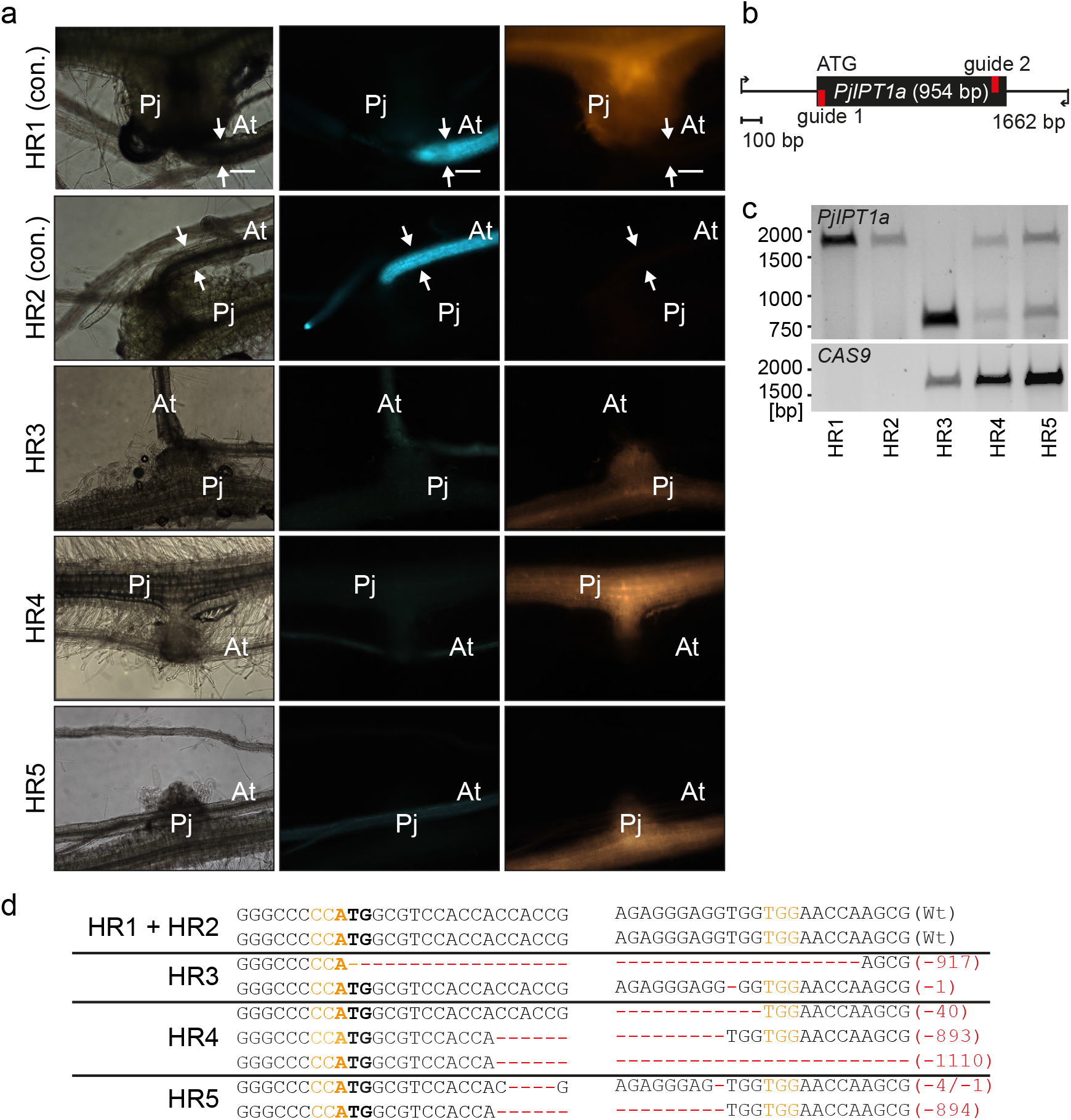
*PjIPT1a* drives host CK responses at the haustorium. (a) Control (con.) haustoria of hairy roots (HR) without the *Cas9/PjIPT1a-gRNA*s cassette (HR1, 2) induced the expression of host CK-response reporter *pARR5:GFP* and hypertrophic growth (arrows) whereas haustoria with the *Cas9/PjIPT1a-gRNA*s cassette (HR3-5) did not. Images were taken at 28 dpi, scale bars = 100 μm. (b) Schematic representation of the *PjIPT1a* gene flanked by the 5’ and 3’ extensions that were amplified in (c). The red colour highlights the target sequences of the guide RNAs. (c) PCR amplicons of the *PjIPT1a* (upper panel) and the *Cas9* genes (lower panel) in *Phtheirospermum* hairy roots were separated on agarose gels and stained with Gel Red. (d) Sequencing results of amplicons shown in c with PAM sequences depicted in orange, the start codon in bold font and Cas9-induced mutations in red show different *PjIPT1a* alleles isolated from indicated hairy roots.

### PjIPT1a and PjIPT1b differ in isopentenyltransferase activity

PjIPT1a and ShIPT1a share a unique triple-proline motif close to the nucleotide-binding site that we only found in IPTa isoforms of parasitic Orobanchaceae species (Fig. **S4**). This raised questions regarding their enzymatic activity. We purified His-tagged PjIPT1a and PjIPT1b from *E. coli* and assayed for isoprenylation of AMP and ATP with DMAPP as the isoprenyl donor and His-tagged GST as negative control (Fig. **4**). When AMP was used as substrate, the total amount of isoprenylated products were similar (factor of 1.3) for PjIPT1a-His and PjIPT1b-His (Fig. **4a**). However, with ATP as substrate, the total amount of isoprenylated products was about four times higher for PjIPT1b-His compared to PjIPT1a-His, indicating higher selectivity for ATP over AMP (Fig. **4b**).

**Figure 4:**
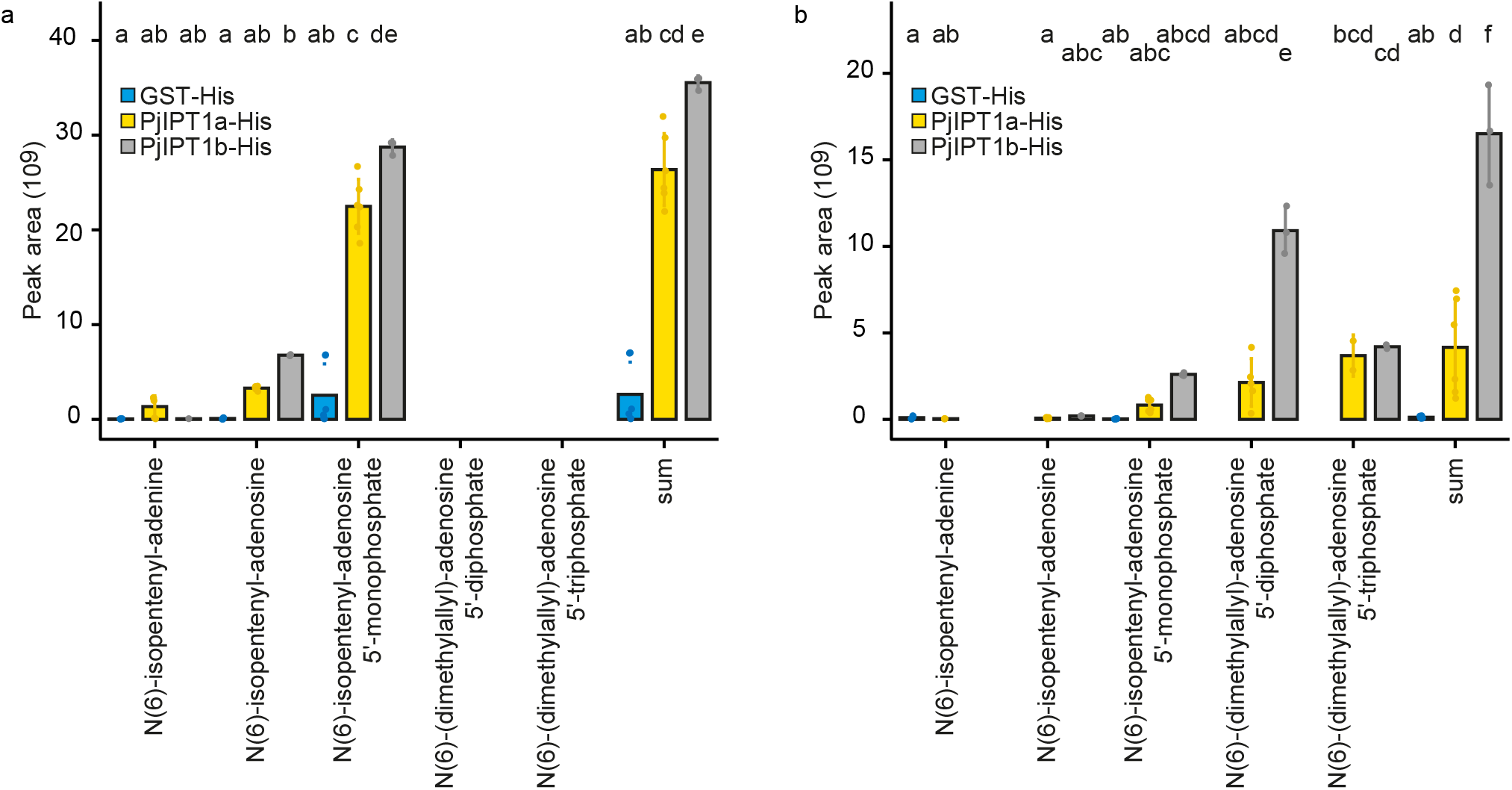
PjIPT1a and PjIPT1b are isopentyltransferases. Plots show average +/− SD (n = 3 – 6) peak areas of extracted-ion chromatograms of individual and cumulative (sum) quantifications of isoprenylated derivatives that were detected in reaction mixtures containing DMAPP, equal amounts of *E. coli* purified GST-His, PjIPT1a-His or PjIPT1b-His, and (a) AMP or (b) ATP as isoprenoid acceptors. Different lower-case letters indicate significant differences according to Tukey HSD with *P* < 0.05. ND, not detected

## Discussion

Here, we studied parasite-induced CK responses in host plants as an example of a neofunctionalized gene function in the Orobanchaceae species *Phtheirospermum japonicum* and *Striga hermonthica*. We found one *IPT* gene, *PjIPT1a*, to be expressed in *Phtheirospermum* haustoria. Reporter gene analysis revealed spatially restricted activity of the *PjIPT1a* promotor in intrusive cells that are in contact with the host vasculature (Fig. **1g-i**). The proximity to the host vasculature is likely sufficient to transfer cytokinins to the host and to trigger CK responses above the haustorium attachment site (Spíchal, 2012). The parasitic tree *Santalum album* also expresses an IPT1 homolog at a similar stage of infections; however, the exact location of *SaIPT1* expressing cells is not known (Zhang *et al.*, 2015).

Unlike in *Santalum album* and many other parasitic plants that are not amenable to reverse genetic manipulation, we were able to use CRISPR/Cas9-based loss-of-function analysis to demonstrate that *PjIPT1a* adopted new parasitism-related functions in *Phtheirospermum*. While RNAi-based knock-down approaches have largely been replaced by the CRISPR/Cas9 technology, RNAi remained, until now, the only strategy to study parasitic plant genes (Bandaranayake *et al.*, 2010; Alakonya *et al.*, 2012; Wakatake *et al.*, 2020). Here, we report the first successful application of the CRISPR/Cas9 technology for studying loss-of-function phenotypes in parasitic plants. CRISPR/Cas9 efficiently introduced deletions in *PjIPT1a* resulting in the loss of up to 98% of the coding sequence, or in the disruption of its open reading frame. In individual hairy roots, we found more than one, in one instance more than two, mutated alleles of *PjIPT1a*, a result of the occasionally chimeric nature of *Agrobacterium rhizogenes-*induced hairy roots (Wakatake *et al.*, 2018). All *Phtheirospermum* roots with non-functional *PjIPT1a* genes failed to induce host CK responses (Fig. **3**).

Our phylogenetic analysis of *IPTs* showed that the duplication event producing *PjIPT1a/ShIPT1a* and *PjIPT1b/ShIPT1b* occurred in a common non-parasitic ancestor of *P. japonicum* and *S. hermonthica* (Fig. **2a**). Surprisingly, *PjIPT1a* and *ShIPT1a*, which were both expressed during infections, were more similar to SrIPT1 from *Streptocarpus rexii* than PjIPT1b and ShIPT1b (Fig. **2a**). The high similarity between parasitism-related and conventional IPTs suggests that neofunctionalization of *PjIPT1a* mainly occurred at the level of its expression profile. Enzymatic constraints may have limited divergence of the gene body and the enzymatic functions of PjIPT1a and b are indeed comparable. However, preference for ATP was reduced in PjIPT1a; its preference for the energetically less costly AMP as isopentenyl donor may reflect an adaptation to the resource-limited parasitic lifestyle that is induced under nutrient-deficient conditions (Spallek *et al.*, 2017). A notable exception to the similarity of PjIPT1a and ShIPT1a to SrIPT1 is a characteristic triple-proline-motif that is only found in parasitism-related IPTs (Fig. **S4**).

Another unexpected feature of PjIPT1a is its nuclear-cytoplasmic localization (Fig. **2b,c**), which it shares with PjIPT1b (Fig. **2d**) and likely also with ShIPT1a and ShIPT1b (Table **S3**). *Arabidopsis* AtIPT1 and AtIPT3 feature functional N-terminal CTPs (Kasahara *et al.*, 2004). However, while AtIPT1 is located in plastids, AtIPT3 is dually targeted, either to the chloroplast, or to the nucleus/cytoplasm in a farnesylation-dependent manner (Galichet *et al.*, 2008). We can exclude a similar mechanism for PjIPT1a, as its predicted CTP appeared to be non-functional, as it was insufficient for targeting GFP to plastids (Fig. **2c**). The relevance of the nuclear-cytoplasmic localization is not clear, but given that it is shared between IPTs that are upregulated during infection and others that are not, a functional link to parasitism seems unlikely.

In conclusion, targeted knock-outs of *PjIPT1a* and its expression in intrusive cells of the haustorium revealed an essential role of PjIPT1a for *Phtheirospermum*-induced CK responses in *Arabidopsis.* The *PjIPT1a* gene likely originated from a non-parasitic ancestral gene through gene duplication followed by neofunctionalization. PjIPT1a has homologs in other parasitic plants. Targeting of neofunctionalized genes in parasitic plants may allow for specific control of parasitic weeds leaving host plants unharmed.

## Acknowledgements

This work supported by the German Research Foundation (DFG, 424122841) to T.S and MEXT KAKENHI JP20H05909 to S.Y. and K.S. The authors thank Ursula Glück-Behrens and Ute Bertsche for technical support.

## Author Contribution

T.S., A.S., and K.S supervised the research. A.G. and I.B. analysed the subcellular location of proteins, purified IPTs from *E. coli* and conducted the *in vitro* assays. J.P. measured the IPT *in vitro* assays compounds by HPLC-MS/MS. T.S. performed the remaining experiments. A.S., P.S., and T.S. analysed data. Y.S. and K.S. provided crucial non-published data. T.S. and A.S. wrote the paper.

## Data Availability

Table **S1** contains accession numbers of genes and proteins used in this study. Gene sequences for *PjIPT1a* and *PjIPT1b* were deposited in the NCBI Gene Bank under the accession numbers MZ441091 and MZ441090, respectively. Previously published sequencing data is available from at the Dryad database (https://doi.org/10.5061/dryad.vt4b8gtpt) and at the National Center for Biotechnology Information, NCBI, (https://www.ncbi.nlm.nih.gov/sra/DRX219041[accn] for *Phtheirospermum japonicum* and https://doi.org/10.5061/dryad.53t3574 for *Striga hermonthica*).

## Notes

### Competing Interest Statement

The authors have declared no competing interest.

