## Supplementary material for "The *Phtheirospermum japonicum* isopentenyltransferase PjIPT1a regulates host cytokinin responses in *Arabidopsis*": Fig. S1

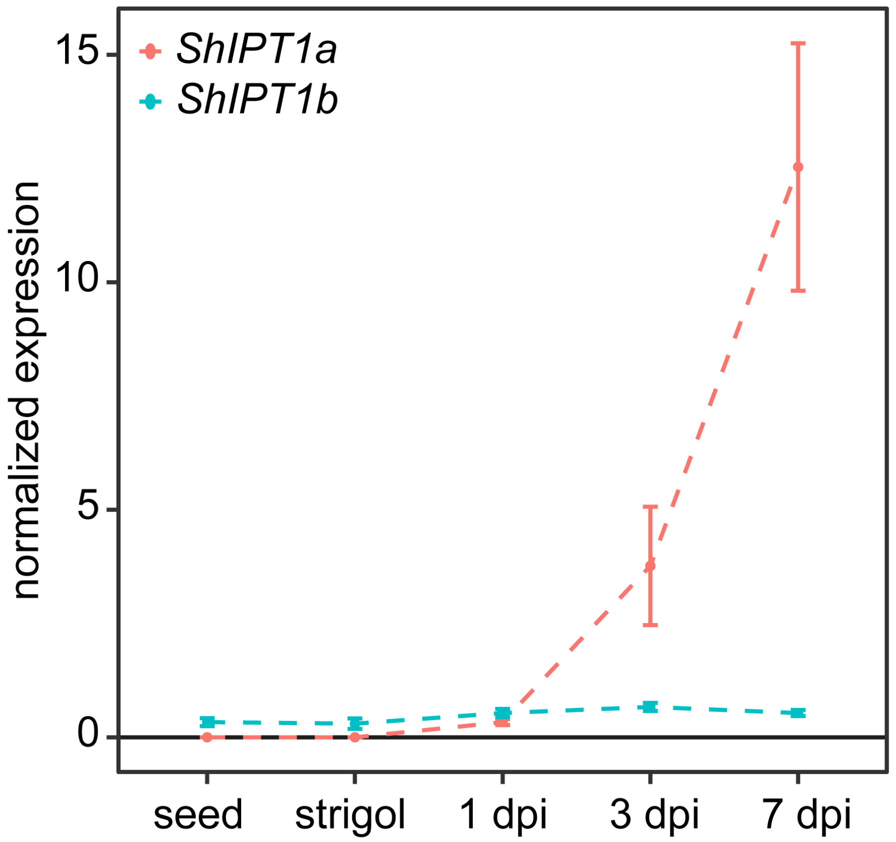


**Supplement Figure S1: *ShIPT1a* but not *ShIPT1b* is up-regulated in *Striga* during rice infections** Mean +/- 1SE normalized expression of *ShIPT1a* (red) and *ShIPT1b* (blue) along different time points of infections. RNA-Sequencing data were published in Yoshida *et al.*, 2019.
