## Supplementary material for "The *Phtheirospermum japonicum* isopentenyltransferase PjIPT1a regulates host cytokinin responses in *Arabidopsis*": Fig. S2

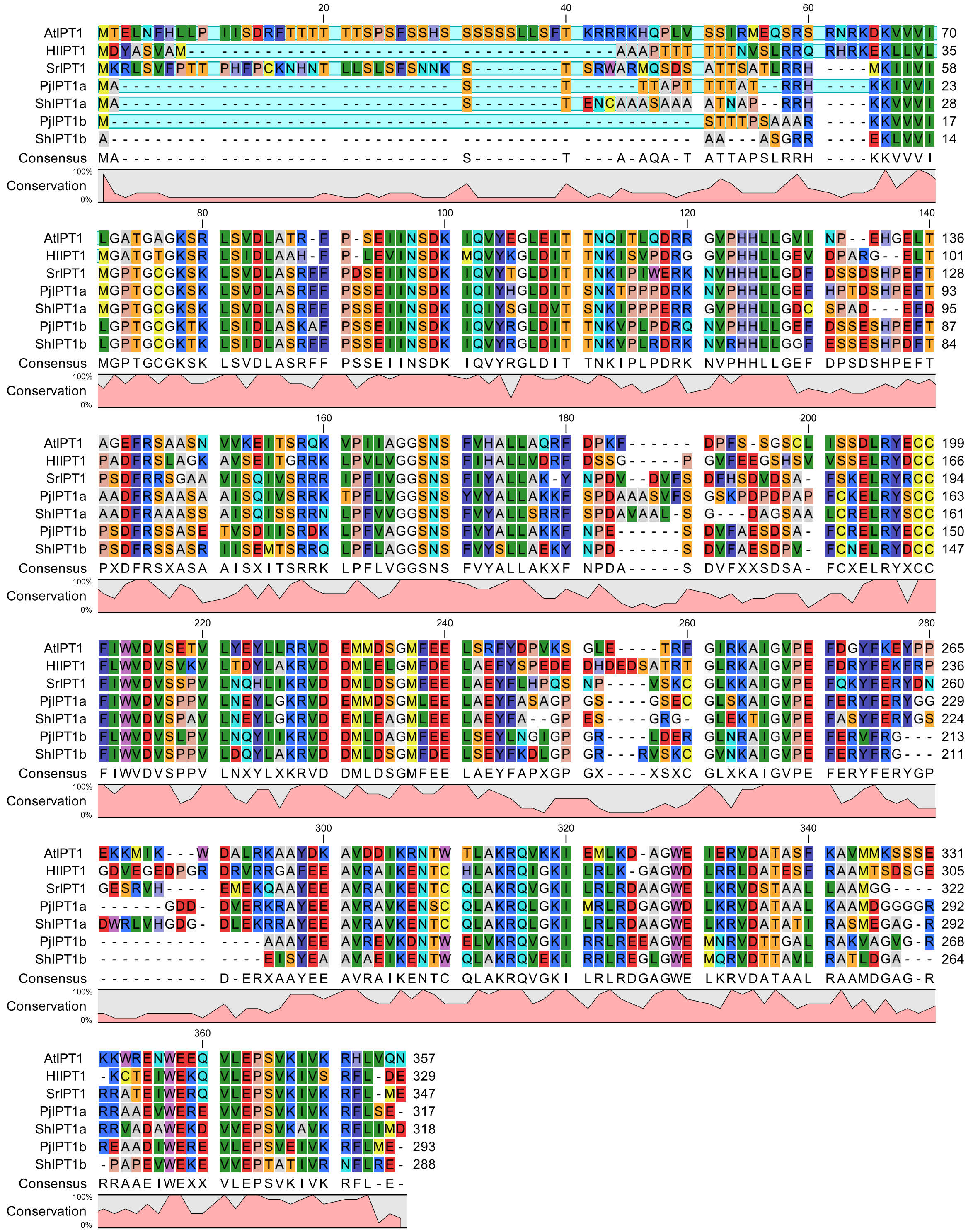


**Supplement Figure S2: Alignments of IPT1 proteins.** Selected IPT1 family proteins were aligned and the predicted chloroplast transit peptides are highlighted in blue.
