## Supplementary material for "The *Phtheirospermum japonicum* isopentenyltransferase PjIPT1a regulates host cytokinin responses in *Arabidopsis*": Fig. S3

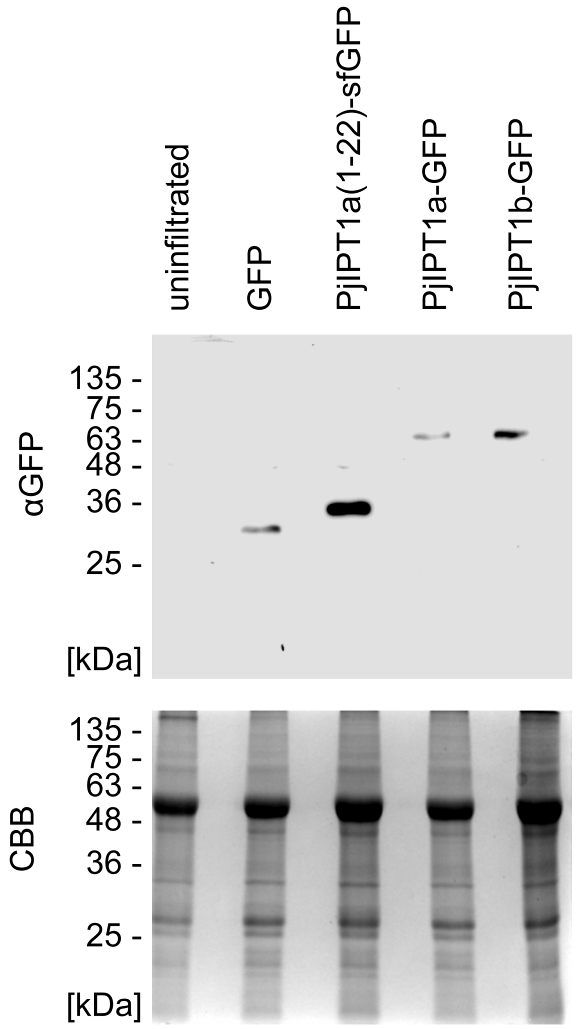


**Supplement Figure 3: Western blot of IPTs used for *N. benthamiana* subcellular localization assays.** Polyclonal anti-GFP antibody (# A-11122, Thermo Fisher Scientific) was used to detect GFP-fusions in *N. benthamiana* total extracts. Coomassie Brilliant Blue (CBB)-stained SDS-PAGE of an equal amount of samples loaded for Western-Blotting indicate comparable loading.
