## Supplementary material for "The *Phtheirospermum japonicum* isopentenyltransferase PjIPT1a regulates host cytokinin responses in *Arabidopsis*": Fig. S4

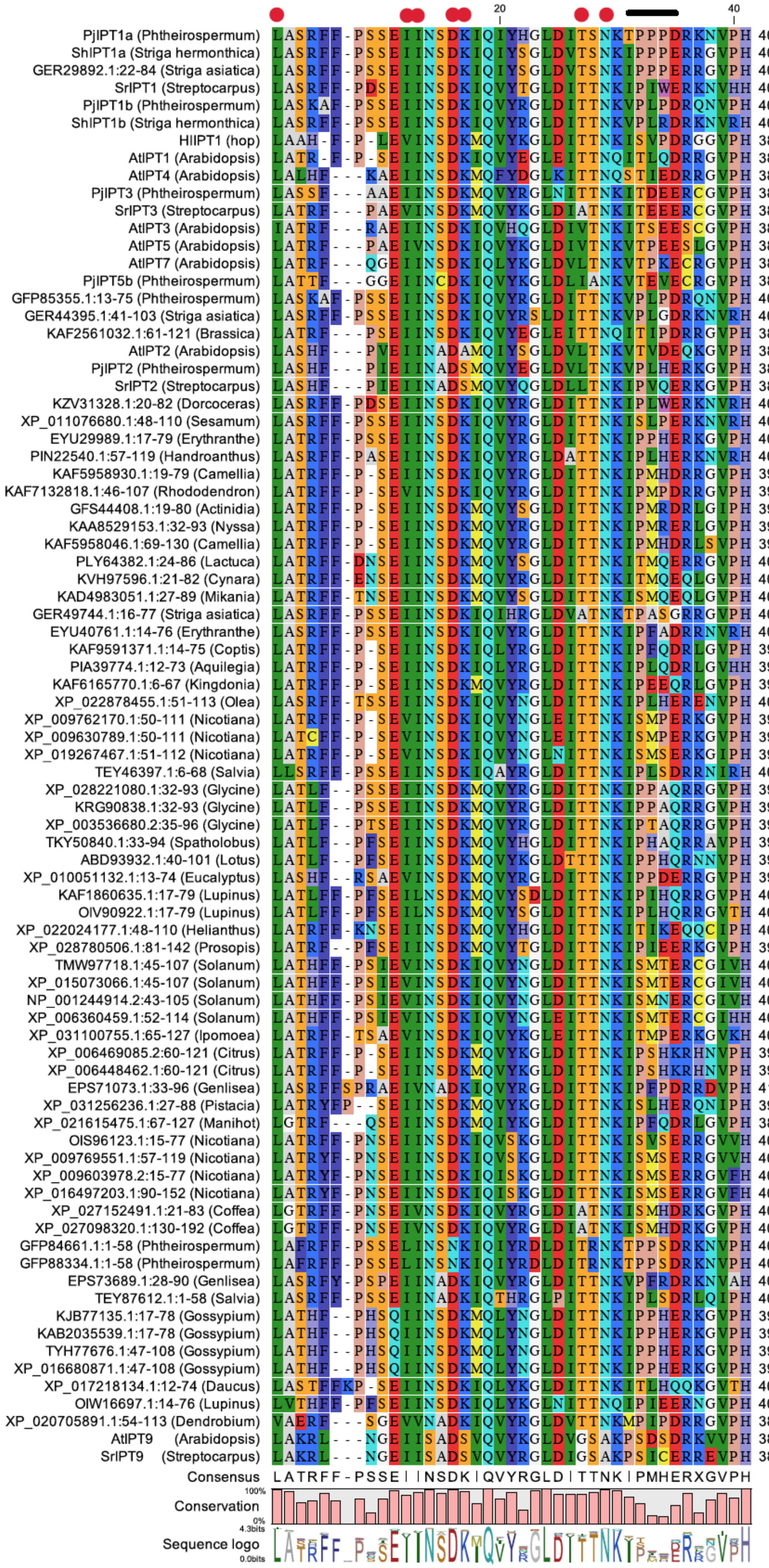
20
40
PjIPT1a (Phtheirospermum) LASRFF-PSSEIINSDKIQIYHGLDITSNKTPPPDRKNVPH 40
ShIPT1a (Striga hermonthica) LASRFF-PSSEIINSDKIQIYSGLDVTSNKIPPPERRGVPH 40
GER29892.1:22-84 (Striga asiatica) LASRFF-PSSEIINSDKIQIYSGLDVTSNKIPPPERRGVPH 40
SrIPT1 (Streptocarpus) LASRFF-PDSEIINSDKIQVYTGLDITTNKIPIWERKNVHH 40
PjIPT1b (Phtheirospermum) LASKAF-PSSEIINSDKIQVYRGLDITTNKVPLPDRQNVPH 40
ShIPT1b (Striga hermonthica) LASRFF-PSSEIINSDKIQVYRGLDITTNKVPLRDRKNVRH 40
HlIPT1 (hop) LAAH-F-P-LEVINSDKMQVYKGLDITTNKISVPDRGGVPH 38
AtIPT1 (Arabidopsis) LATR-F-P-SEIINSDKIQVYEGLEITTNQITLQDRRGVPH 38
AtIPT4 (Arabidopsis) LALHF---KAEIINSDKMQFYDGLKITTNQSTIEDRRGVPH 38
PjIPT3 (Phtheirospermum) LASSF---AAEIINSDKMQVYRGLNITTNKITDEERCGVPH 38
SrIPT3 (Streptocarpus) LATRF---PAEVINSDKMQVYKGLDIATNKITEEERCGVPH 38
AtIPT3 (Arabidopsis) IATRF---RAEIINSDKIQVHQGLDIVTNKITSEESCGVPH 38
AtIPT5 (Arabidopsis) LATRF---PAEIVNSDKIQVYKGLDIVTNKVTPEESLGVPH 38
AtIPT7 (Arabidopsis) LATRF---QGEIINSDKIQLYKGLDVLTNKVTPKECRGVPH 38
PjIPT5b (Phtheirospermum) LATTF---GGEIINCDKIQVYKGLDLIANKVTEVECRGVPH 38
GFP85355.1:13-75 (Phtheirospermum) LASKAF-PSSEIINSDKIQVYRGLDITTNKVPLPDRQNVPH 40
GER44395.1:41-103 (Striga asiatica) LASRFF-PSSEIINSDKIQVYRSLDITTNKVPLGDRKNVRH 40
KAF2561032.1:61-121 (Brassica) LATRF---PSEIINSDKIQVYEGLEITTNQITIPDRRGVPH 38
AtIPT2 (Arabidopsis) LASHF---PVEIINADAMQIYSGLDVLTNKVTVDEQKGVPH 38
PjIPT2 (Phtheirospermum) LASHF---PIEIINADSMQVYEGLDVLTNKVPLHERKGVPH 38
SrIPT2 (Streptocarpus) LASHF---PIEIINADSMQVYQGLDLLTNKIPVQERKGVPH 38
KZV31328.1:20-82 (Dorcoceras) LASRFF-PDSEIINSDKIQVYRGLDITTNKIPLWERKNVRH 40
XP_011076680.1:48-110 (Sesamum) LASRFF-PSSEIINSDKIQVYRGLDITTNKISLPERKNVRH 40
EYU29989.1:17-79 (Erythranthe) LATRFF-PSSEIINSDKIQVYRGLDITTNKIPPHERKGVPH 40
PIN22540.1:57-119 (Handroanthus) LASRFF-PASEIINSDKIQVYRGLDATTNKIPLHERKNVRH 40
KAF5958930.1:19-79 (Camellia) LATRFF-P-SEIINSDKIQVYRGLDITTNKIPMHDRRGVPH 39
KAF7132818.1:46-107 (Rhododendron) LATRFF-P-SEVINSDKIQVYRGLDITTNKIPMPDRRGVPH 39
GFS44408.1:19-80 (Actinidia) LATRFF-P-SEIINSDKMQVYSGLDITTNKIPMRDRLGIPH 39
KAA8529153.1:32-93 (Nyssa) LATRFF-P-SEIINSDKMQVYRGLDITTNKIPMRERLGVPH 39
KAF5958046.1:69-130 (Camellia) LATRFF-P-SEIINSDKMQVYRGLDITTNKIPMHDRLSVPH 39
PLY64382.1:24-86 (Lactuca) LATRFF-DNSEIINSDKMQVYSGLDITTNKITMQERRGVPH 40
KVH97596.1:21-82 (Cynara) LATRFF-ENSEIINSDKMQVYRGLDITTNKITMQEQLGVPH 40
KAD4983051.1:27-89 (Mikania) LATRFF-TNSEIINSDKMQVYSGLDITTNKISMQEQLGVPH 40
GER49744.1:16-77 (Striga asiatica) LASRFF-PSSEIINSDKIQIHRGLDVATNKTPASGRRGVPH 40
EYU40761.1:14-76 (Erythranthe) LASRFF-PSSEIINSDKIQVYRGLDITTNKIPFADRRNVRH 40
KAF9591371.1:14-75 (Coptis) LATRFF-P-SEIINSDKIQLYRGLDITTNKIPFQDRLGVPH 39
PIA39774.1:12-73 (Aquilegia) LATRFF-P-SEIINSDKIQLYRGLDITTNKIPLQDRLGVHH 39
KAF6165770.1:6-67 (Kingdonia) LATRFF-P-SEIINSDKMQVYRGLDITTNKIPEEQRLGVPH 39
XP_022878455.1:51-113 (Olea) LASRFF-TSSEIINSDKIQVYNGLDITTNKIPLHERENVPH 40
XP_009762170.1:50-111 (Nicotiana) LATRFF-P-SEVINSDKIQVYNGLEITTNKISMPERKGVPH 39
XP_009630789.1:50-111 (Nicotiana) LATCFF-P-SEVINSDKIQVYNGLEITTNKISMPERKGVPH 39
XP_019267467.1:51-112 (Nicotiana) LATRFF-P-SEVINSDKIQVYNGLNITTNKISMSERKGIPH 39
TEY46397.1:6-68 (Salvia) LLSRFF-PSSEIINSDKIQAYRGLDITTNKIPLSDRRNIRH 40
XP_028221080.1:32-93 (Glycine) LATLF--PSSEIINSDKMQVYRGLDITTNKIPPAQRRGVPH 39
KRG90838.1:32-93 (Glycine) LATLF--PSSEIINSDKMQVYRGLDITTNKIPPAQRRGVPH 39
XP_003536680.2:35-96 (Glycine) LATLF--PTSEIINSDKMQVYRGLDITTNKIPTAQRRGVPH 39
TKY50840.1:33-94 (Spatholobus) LATLF--PFSEIINSDKMQVYHGLDITTNKIPHAQRRAVPH 39
ABD93932.1:40-101 (Lotus) LATLF--PFSEIINSDKMQVYKGLDTTTNKIPPHQRNNVPH 39
XP_010051132.1:13-74 (Eucalyptus) LASHF--RSAEVINSDKMQVYRGLDITTNKIPPDERRGVPH 39
KAF1860635.1:17-79 (Lupinus) LATLFF-PFSEILNSDKMQVYSDLDITTNKIPIHQRRGVPH 40
OIV90922.1:17-79 (Lupinus) LATLFF-PFSEILNSDKMQVYSGLDITTNKIPLHQRRGVTH 40
XP_022024177.1:48-110 (Helianthus) LATRFF-KNSEIINSDKMQVYHGLDITTNKITIKEQQCIPH 40
XP_028780506.1:81-142 (Prosopis) LATRF--PFSEIINSDKMQVYTGLDITTNKIPIEERKGVPH 39
TMW97718.1:45-107 (Solanum) LATHFF-PSIEVINSDKIQVYNGLDITTNKISMTERCGIVH 40
XP_015073066.1:45-107 (Solanum) LATHFF-PSIEVINSDKIQVYNGLDITTNKISMTERCGIVH 40
NP_001244914.2:43-105 (Solanum) LATHFF-PSIEVINSDKIQVYNGLDITTNKISMNERCGIVH 40
XP_006360459.1:52-114 (Solanum) LATHFF-PSIEVINSDKIQVYNGLEITTNKISMTERCGIHH 40
XP_031100755.1:65-127 (Ipomoea) LATRFF-TSAEVINSDKIQVYKGLEITTNKITMPERKGVKH 40
XP_006469085.2:60-121 (Citrus) LATRFF-P-SEIINSDKMQVYKGLDITTNKIPSHKRHNVPH 39
XP_006448462.1:60-121 (Citrus) LATRFF-P-SEIINSDKMQVYKGLDITTNKIPSHKRHNVPH 39
EPS71073.1:33-96 (Genlisea) LASRFFSPRAEIVNADKIQVYRGLDITTNKIPFPDRRDVPH 41
XP_031256236.1:27-88 (Pistacia) LATRYFP--SEIINSDKMQVYKGLDITTNKISLHERQNIPH 39
XP_021615475.1:67-127 (Manihot) LGTRF---QSEIINSDKMQIYKGLDITTNKIPFQDRLGVPH 38
OIS96123.1:15-77 (Nicotiana) LATRFF-PNSEIINSDKIQVSKGLDITTNKISVSERRGVVH 40
XP_009769551.1:57-119 (Nicotiana) LATRYF-PNSEIINSDKIQVSKGLDITTNKISMSERRGVVH 40
XP_009603978.2:15-77 (Nicotiana) LATRYF-PNSEIINSDKIQISKGLDITTNKISMSERRGVFH 40
XP_016497203.1:90-152 (Nicotiana) LATRYF-PNSEIINSDKIQISKGLDITTNKISMSERRGVFH 40
XP_027152491.1:21-83 (Coffea) LGTRFF-PNSEIVNSDKIQVYRGLDIATNKISMHDRKGVPH 40
XP_027098320.1:130-192 (Coffea) LGTRFF-PNSEIVNSDKIQVYRGLDIATNKISMHDRKGVPH 40
GFP84661.1:1-58 (Phtheirospermum) LAFRFF-PSSELINSNKIQIYRDLDITRNKTPPSDRKNVPH 40
GFP88334.1:1-58 (Phtheirospermum) LAFRFF-PSSELINSNKIQIYRDLDITRNKTPPSDRKNVPH 40
EPS73689.1:28-90 (Genlisea) LASRFY-PSPEIINADKIQVYRGLDITTNKVPFRDRKNVAH 40
TEY87612.1:1-58 (Salvia) LASRFF-PSSEIINADKIQTHRGLPITTNKIPLSDRLQIPH 40
KJB77135.1:17-78 (Gossypium) LATHF--PHSQIINSDKMQLYNGLDITTNKIPPHERKGVPH 39
KAB2035539.1:17-78 (Gossypium) LATHF--PHSQIINSDKMQLYNGLDITTNKIPPHERKGVPH 39
TYH77676.1:47-108 (Gossypium) LATHF--PHSQIINSDKMQLYNGLDITTNKIPPHERKGVPH 39
XP_016680871.1:47-108 (Gossypium) LATHF--PHSQIINSDKMQLYNGLDITTNKIPPHERKGVPH 39
XP_017218134.1:12-74 (Daucus) LASTFFKP-SEIINSDKIQLYKGLDITTNKITLHQQKGVTH 40
OIW16697.1:14-76 (Lupinus) LVTHFF-PFSEIINSDKIQVYKGLNITTNQIPIEERNGVPH 40
XP_020705891.1:54-113 (Dendrobium) VAERF---SGEVVNADKIQVYRGLDVTTNKMPIPDRRGVPH 38
AtIPT9 (Arabidopsis) LAKRL---NGEIISADSVQVYKGLDVGSAKPSDSDRKVVPH 38
SrIPT9 (Streptocarpus) LAKRL---NGEIISADSVQVYRGLDIGSAKPSICERREVPH 38
Consensus LATRFF-PSSEIINSDKIQVYRGLDITTNKIPMHERXGVPH
100%
Conservation
0%
4.3bits
Sequence logo
0.0bits

**Supplement Figure S4: Alignment of the IPT nucleotide-binding domain across plant species.** Red dots indicate residues that participate in substrate binding, the black line marks the position of the triple-proline motif
